# Action-outcome based flexible behavior requires medial prefrontal cortex lead and its enhanced functional connectivity with dorsomedial striatum

**DOI:** 10.1101/2023.12.12.571355

**Authors:** Áron Kőszeghy, Wei Xu, Mingshan Liu, Peiheng Lu, Long Wan, Peggy Series, Jian Gan

**Author notes:** Corresponding author: Dr. Jian Gan, UK Dementia Research Institute Centre for Discovery Brain Sciences The University of Edinburgh Chancellor’s Building, 49 Little France Crescent Edinburgh, EH16 4SB United Kingdom. Present address: Institute of Neurochemistry, Medical University of Innsbruck (Áron Kőszeghy). These authors contributed equally to this work (Áron Kőszeghy, Wei Xu).

## Abstract

Cognitive flexibility plays a key role in ensuring an individual’s survival, and its deficit is a key symptom in many mental conditions and neurodegenerative diseases. The prefrontal cortex and striatum are both essential to cognitive flexibility. However, how the prefrontal cortex and striatum communicate with each other to enable flexible decision-making is not well understood. Competing theories are raised, debating on which structure among these two leads the role in detecting and representing the new circumstances for a change, giving largely opposing predictions on neural activities in the prefrontal cortex and striatum during flexible behavior.

To address this question, we trained head-restrained mice to perform an action-outcome based dynamic foraging task and simultaneously recorded single-neuron activities in the medial prefrontal cortex (mPFC) and dorsomedial striatum (DMS). In this task, the animal chooses one of two actions to obtain reward. The animal is guided only by previous reward outcomes. We report that mPFC but not DMS activity stores information about prior reward history. A large fraction of both mPFC and DMS neurons’ activity represents the difference in reward probability between two alternative options, namely the perceived reward probability difference (PRPD), a key decision variable that prescribes which subsequent choice to make. We find that mPFC neural activities track the change of PRPD earlier and faster than those in the DMS, and functional connectivity between mPFC and DMS increases with reducing overall reward proportion.

## Introduction

The ability to adapt one’s actions and choices to mitigate risks and optimize rewards in response to changes in the environment plays a crucial role in survival. The prefrontal cortex (PFC) is a critical brain region for flexible decision making^1-5^ by monitoring behavioral performance and executing top-down control^6-9^. However, PFC does not operate in isolation. Notably, the striatum, which serves as the input nucleus of the basal ganglia (BG), plays a crucial role in enabling all forms of flexible goal-directed behavior^10,11^. More specifically, the medial prefrontal cortex (mPFC) and dorsomedial striatum (DMS) are implicated in action-outcome based flexible behaviors, where the choice of action is based on the evaluation of outcomes from previous actions on a trial-by-trial basis^2,10-14^.

mPFC and DMS have been typically studied separately, involving different preparations and tasks^15-18^. How these two regions communicate with each other in enabling action-outcome based flexible behavior is not well understood. It is also not clear how behavioral representations in these two brain regions evolve and influence one another during flexible behavior. Two competing hypotheses have been proposed. The first posits that the striatum is at the center of learning the new rule for a change as ‘context detector’, due to its nature as an anatomical ‘hub’, connecting neocortical and sub-cortical nuclei, and the high convergence nature of medium spiny neurons, which are ideally suitable for reinforcement learning-related plasticity^19,20^. This model suggests that prefrontal activities could be ‘trained’ by the striatum, predicting that DMS activities for detecting ‘a change of context’ will lead mPFC activities during flexible behavior. The second theory suggests that the prefrontal cortex is responsible for learning the new situation and the striatum subsequently consolidates newly learned routines into familiar habits^21,22^. This model predicts the opposite that mPFC will lead DMS.

To tackle this question, we trained head-restrained mice to perform an action-outcome association based dynamic foraging task whilst simultaneously recording single-neuron activities in the mPFC and DMS using high-channel-count silicon probes. This task is purely based on action-outcome history. No sensory cue instructs responses for the favorable option. A reinforcement learning (Q-learning) model was used to quantitatively describe the animal’s performance and predict its choice behavior reliably^23,24^. We discovered that a large fraction of both mPFC and DMS neurons represented the difference in respective reward history from the two alternative options, namely ‘perceived reward probability difference (PRPD), a key decision variable that biases which subsequent choice to make. In addition, the mPFC neural activities tracked the change of PRPD earlier and faster than those in the DMS. Further analysis showed that in a large proportion of putative monosynaptic mPFC-DMS neuron pairs, one or both members encoded PRPD. Furthermore, the spike-to-spike transmission efficacy between these pairs was largely negatively correlated with the overall reward proportion.

Taken together, our data suggest the mPFC is likely to be the first responder and the source of encoding the dynamic difference in reward probabilities. Computed decision signals are then transmitted via the mPFC-to-DMS projections to bias the animal’s choice. In addition, the strength of information transfer between the mPFC and DMS, as measured by the efficacy of actional potential transmission between them, increases when there is a strong need to change behavioral strategy. Our results support a framework where the mPFC tracks the change of decision variables faster and earlier than the DMS, and the computed decision signal is sent downstream to the DMS to bias behavioral engagement to advantageous strategies, which in turn can be implemented by subordinate hierarchical cortico-BG circuits.

## Results

### Dynamic foraging in head-restrained mice enables the study of prefronto-striatal correlates of action-outcome based flexible behavior

To investigate prefronto-striatal neural dynamics underlying goal-directed flexible behavior, we first developed a steering wheel choice device-based^25^ probabilistic dynamic foraging paradigm^15^ in head-restrained mice (see also methods). In this task, mice were given two alternative options (moving the wheel left or right) to obtain a drop of liquid reward (Fig. 1A). The temporal structure of a trial is illustrated in Fig.1B (Top). In a single trial, the mean wheel angular speed which reflects the animal turning the wheel, and the distribution of the timing of reward delivery over the course of a trial are plotted in Fig.1B (Bottom, calculated from all sessions). The time of reward delivery peaked around 0.36s after trial initiation (at t=0), and non-rewarded trials have a longer tail in the wheel’s angular speed – likely due to the animal persisting in turning the wheel when an expected reward was not delivered. The probabilities of reward on turning the wheel to the left or right were not equal, with one side always having a much higher reward probability than the other. The mean number of trials per behavioral session was 396.2 ± 90.3 trials, and the mean response time was 0.92 ± 0.02s (median: 0.39s). A representative session is shown in Fig.1C. The two sides’ reward probabilities stayed constant for a variable number of trials, which we will refer to as a block of trials (mean block duration = 116.1655 ± 52.3498 trials, 5 animals, 29 sessions). After the end of each block, the high and low probability sides were reversed, unannounced to the animal. This reversal occurs several times in a session (mean number of blocks per session = 3.8 ± 1.1, 5 animals, 29 sessions).

**Figure 1.**
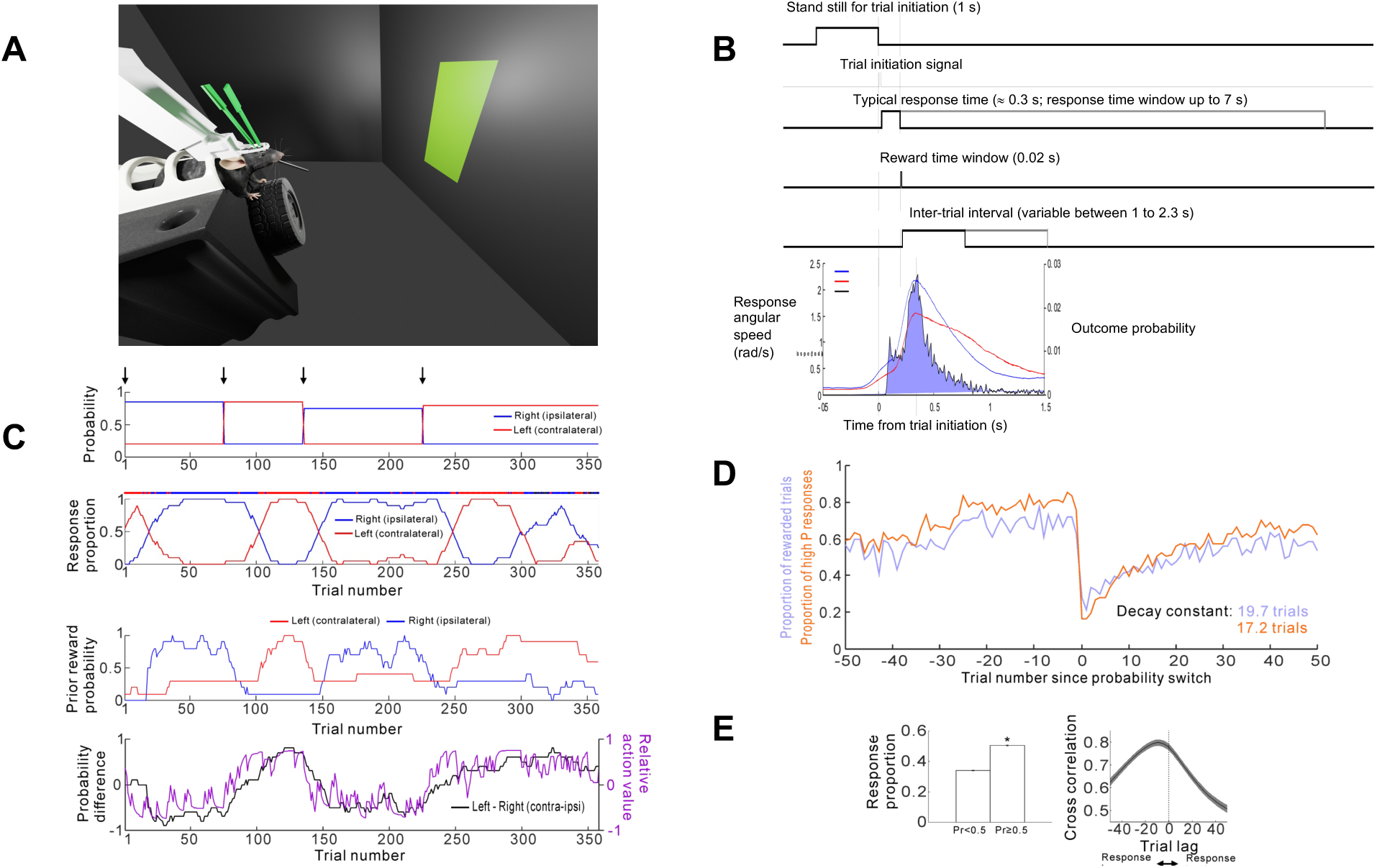
Quantitative description of action-outcome based dynamic foraging task in head-restrained mice. **A.** To study goal-directed flexible behavior, we developed a modified version of the value-based probabilistic dynamic foraging paradigm^15^, in which head-fixed animals express their choice by turning a steering wheel left or right to obtain liquid rewards^25^. **B**. A schematic diagram shows the temporal structure of the probabilistic dynamic foraging task. In a single trial, animals were required to withhold wheel rotation for 1 second to initiate a trial, where they had 7 s to respond (mean response time: 0.92 ± 0.02s, median: 0.39s). Responses were followed by a variable duration inter-trial-interval (ITI). **C**. Animal choices and reward probabilities from an example session. Top: Set reward probabilities for the left and right. Upper middle: Proportion of left and right responses made by the animal. Lower middle: Perceived left and right reward probability. Bottom: Perceived reward probability difference of the left and right options (PRPD, black curve) and relative value (ΔQ, Q_left_ – Q_right_) computed from Q-learning model (magenta curve) in this session. **D**. Proportion of rewarded trials (blue) and advantageous responses (orange) aligned by switches in 79 block transitions of the 29 recording sessions. **E**. Left: comparison of mean proportion of responses to the side with <0.5 reward probability and >0.5 reward probability (proportions calculated for each session and then averaged. Error bars indicate standard error, * indicates P<0.05, paired t-test). Right: Mean cross correlation between set probability for a given side and response proportions to that same side. Shaded regions indicate standard error.

In order to measure the reward history, and therefore a surrogate of the animal’s perception of current reward probability based on past events, we calculated for each side the proportion of rewards received in the last ten trials for responses to that side (perceived/prior reward probability, PRP). The PRP values from the same example session are plotted (Fig.1C, bottom upper). As a measure of the perceived reward probability difference (PRPD) between the two sides (and therefore a gauge of the animal’s internal drive towards the advantageous side), we subtracted the PRP of the ipsilateral side from the contralateral side (relative to the hemisphere being recorded) – example plotted in Fig.1C (bottom lower, black trace), showing this metric to modulate similarly with response side choice (Fig.1C, middle, blue trace). We also quantitatively described mice foraging behavior in our task using a generative reinforcement learning model (Q-learning), which has been used in similar dynamic foraging paradigms from earlier studies^15,16^. Consistent with these studies, we find the animal’s choices in our task strongly overlap with the prediction from this Q-learning model (Fig.1C, bottom lower, magenta trace). In addition, the calculated relative values (ΔQ, the difference between the left and right action values, Q_left_ − Q_right_), which have been shown to bias the animal’s choice previously^15^, overlap tightly with the PRPD values in our task (see an example session in Fig.1C, bottom panel), suggesting PRPD is a simplified yet reliable quantitative measure of reward history of the animal and can be used as a proxy of relative value (Δ1Q).

In our task, on average animals chose the high probability side significantly more than the low probability side (Fig.1E, left). It took several trials after a probability switch (block transition) for the animal to discover the new advantages side and adapt their strategy to favor the other direction, therefore the proportion of responses to a given side lags the changes in set probability for that side, as shown by their averaged cross-correlogram (Fig.1E, right). In order to chart how quickly an animal adapts to new reward rules after a switch, we looked at task performance before and after block transitions. The overall proportion of rewarded trials and advantageous responses are aligned by switch points (from all trials in all sessions, averaged over trials for each trial) in Fig.1D. Both parameters drop sharply after the switch and only start to increase when the animal starts to adapt to the new advantageous strategy. On average animals learned the change with newly adapted behavior plateaus around 30 trials post-switch. The time constants for the proportion of rewarded trials and the proportion of advantageous responses (therefore the speed of behavioral adaption to the changed reward probabilities) were 19.7 and 17.2 trials respectively.

### Modulation of neural activities during single task trials in the mPFC and DMS

By using two 128-channel silicon probes, we recorded single units and local field potentials from 5 mice for a total of 29 sessions from mPFC and the ipsilateral DMS during the dynamic foraging task. The average coordinates of all 29 sessions were visualized in the PinPoint software (Fig. 2A-F). Post-hoc histology analysis confirmed the locations of the recording electrodes. In total, following clustering,1668 neurons from mPFC and 974 neurons from DMS were recorded during the performance of the task.

**Figure 2.**
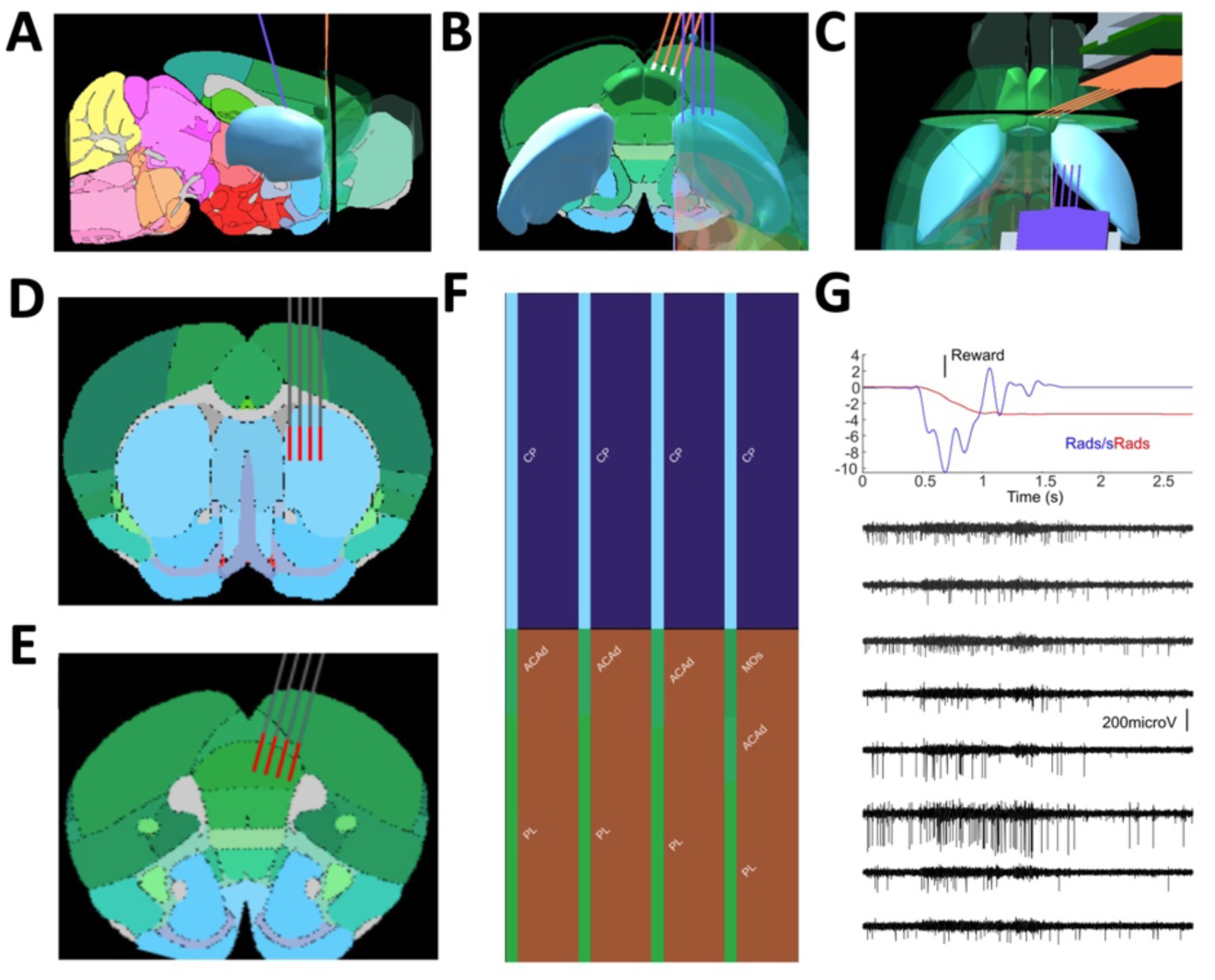
Simultaneous high-density silicon probe recordings in the mPFC and DMS during the dynamic foraging task. **A-C.**Simultaneous dual silicon probe recordings were performed from the medial prefrontal cortex (mPFC) and the dorsomedial striatum (DMS) using 4-shank, 128-channel high-density silicon probes. Using in vivo MRI-morphed Allen mouse common coordinates framework of the PinPoint software panels A to C show sagittal, coronal, and horizontal views of a mouse brain with two silicon probes set to the average coordinates of all recording sessions. 3D contours of the prelimbic cortex and dorsal striatum are shown in green and blue, respectively. **D and E.** The average probe tracks in DMS and mPFC, respectively. Using the in vivo MRI-morphed Allen mouse common coordinates framework of the PinPoint software. **F.** The pinpoint readout of brain regions along the average tracks (CP: caudate putamen; ACAd: dorsal anterior cingulate cortex; PL: prelimbic cortex; MOs: secondary motor cortex). **G.** Example trial showing the angular velocity of the wheel (blue), total angle traversed (red), and high-pass filtered original single-unit recording traces from eight example channels simultaneous multiple in the mPFC.

Previous studies have shown the mPFC units can be modulated by multiple behavioral variables^26^, and often single units can respond to more than one of those, reflecting their high dimensional coding with mixed selectivity^27-29^. To quantitatively measure how spiking modulations are influenced by different task variables in our paradigm, we first examined the spiking patterns of mPFC and DMS neurons during the time course of single trials (time scale of seconds). We then focused on spike modulation during the entire behavioral session on cross-trial level (time scale of mins). For single trial analysis, trials were first separated trials into 3 categories – S - those with trial initiation signals only (i.e. no subsequent response from the animal, no supra-threshold wheel movement), SM - those with signal and wheel movement but no reward, and SMR - those with signal, wheel movement, and reward. Mean mPFC spike firing for these three types of trials are aligned by trial initiation signal and shown in Fig.3A. Signal-only trials elicit a modest increase of firing. Trials with wheel movement and reward elicit a much larger initial increase in spike firing rate. Additionally, non-rewarded trials with wheel movement have a small late increase of spike firing rate – likely to be associated with the small late additional wheel turning when no reward was forthcoming (see Fig.1B bottom). The response profile of DMS neurons for these three types of trials is broadly similar to prefrontal neurons (Fig.3B).

**Figure 3.**
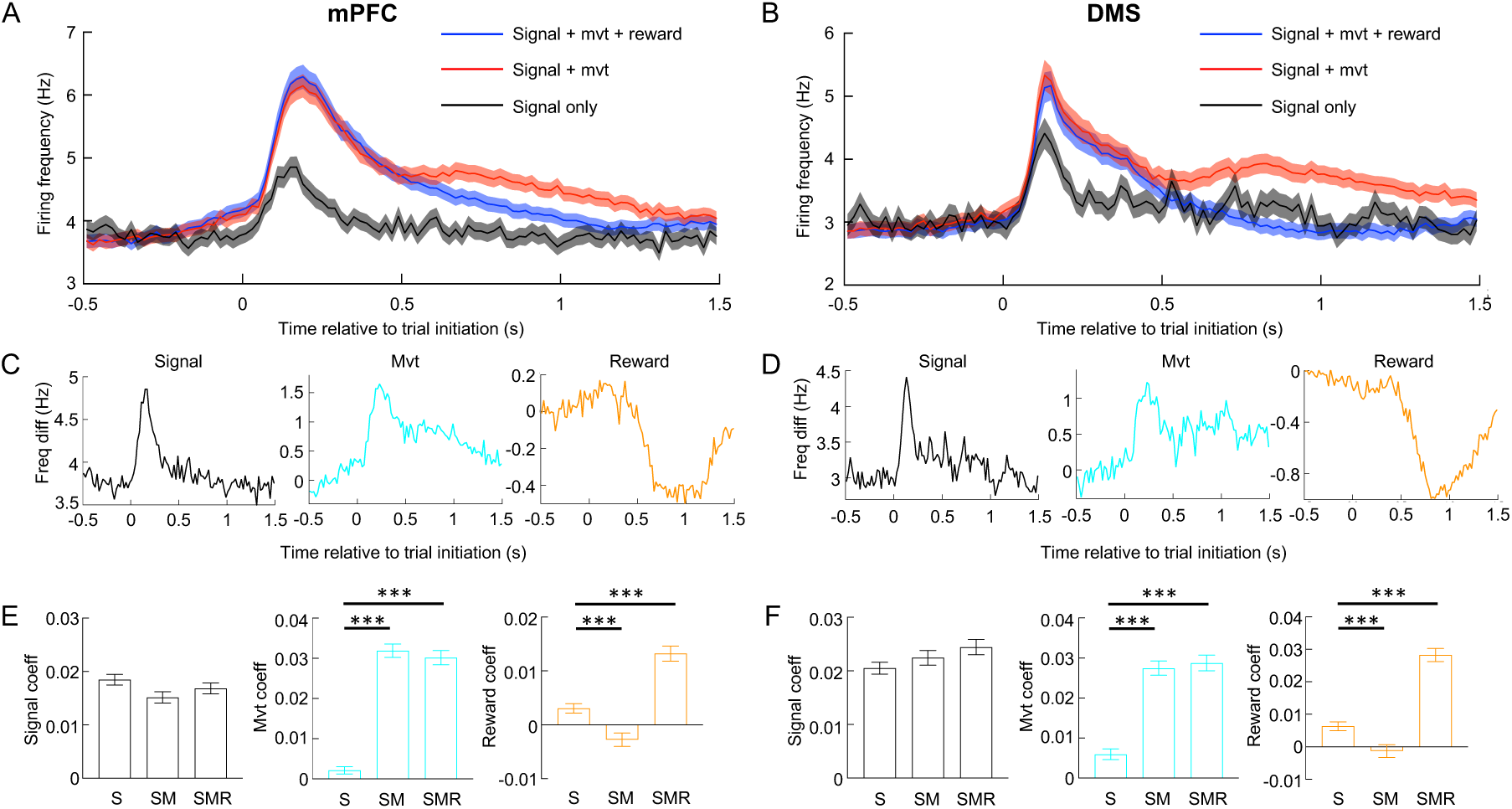
Modulation of neural activities during single task trials in the mPFC and DMS. **A**. Mean mPFC neurons’ spiking frequency is aligned by trial initiation signal for signal-only (S), signal + movement (SM), and signal + movement + reward (SMR) trials over the course of single trials. **B**. Same as A but for DMS neurons. **C**. Signal-related, movement-related, and reward-related regressors were generated to determine the within-trial behavioral modulation of mPFC neurons’ activities using multiple linear regression. **D**. Same as C but for DMS neurons. **E**. Mean regression coefficients for each regressor and each type of trial, compared for S, SM, and SMR trials for the validity of this approach for mPFC neurons. *** represents a statistically significant difference (5% confidence level, one-way ANOVA with post hoc Tukey-Kramer test). **F**. Same analysis for DMS neurons. Error bars and shaded regions indicate standard error of mean (SEM).

Next, we sought to examine how spiking profiles are modulated by the three task modalities namely trial initiation signal, wheel movement, and reward respectively. For trial initiation signal-associated spike firing we used mean spike firing in S trials (Fig.3C left). For wheel movement-associated spike firing we subtracted the mean firing rates of S trials from SM trials (thereby removing the signal component from the spiking profile – Fig.3C middle). For firing profile associated with reward, we subtracted the mean firing rates of SM trials from SMR trials (Fig 3C right). N.B. The later decreasing part (‘tail’) of the reward-related firing profile is likely associated with the prolonged wheel-turning motoric behavior when an expected reward is not delivered. Therefore, we do not claim that this signal is purely reward modulated but simply reward-associated. These three task modality-associated firing modulations were averaged across all neurons for each brain region, scaled to have amplitudes of one, and used as regressors in a multiple linear regression fit to assess the spiking modulation of each individual (also scaled to give unitary amplitudes). The values of the fitted regression coefficients for each regressor were used to gauge the salience of representation for each task modality in the spiking modulation for a particular neuron and under a particular context. To test the validity of this approach, we fitted the three regressors from the mPFC to mPFC neuronal responses separately for S, SM and SMR trials. Since all trials contain a trial-initiation signal the signal regressor’s coefficient was similarly high for all three types of trials (Fig.3E, left). The wheel movement-related coefficient was much lower for trials without wheel movements (Fig.3E, middle) and the reward-related coefficient was much higher in rewarded trials compared to non-rewarded trials (Fig. 3E, right). A similar pattern was found in DMS neurons (Fig. 3F). A 10-fold cross-validation was performed to guard against over-fitting (see methods).

### mPFC but not DMS spiking represents overall reward history information

As the history of prior rewards is the only parameter in this behavioral paradigm that can influence the animals’ choice toward an optimal strategy, it is crucial to examine where and how this information is encoded in the PFC-DMS system. First, for each rewarded trial (i.e. SMR trial), we calculated the proportion of rewarded trials in the prior 10 trials (regardless of chosen side), and separated them into prior reward proportion (PRP) above 0.5 (High PRP) and PRP below 0.5 (Low PRP). Second, we gauged whether the modality-specific regression coefficients were influenced by PRP, therefore reward history information. For mPFC neurons the signal-related regression coefficients between high and low PRPs were similar (Fig.4A, left). The wheel movement-related coefficients were also unchanged (Fig.4A, middle). However, the fitted reward-related coefficient was profoundly smaller for low prior reward trials (Fig.4A, right). In contrast, none of these coefficients were different between high and low PRPs for DMS neurons (Fig.4B). These results suggest that mPFC spiking profile represents both rewards delivered in the current trial as well as rewards delivered in previous trials, a ‘working memory’ of reward history that the animal has experienced, whereas DMS spiking seems indifferent to prior reward experience.

**Figure 4.**
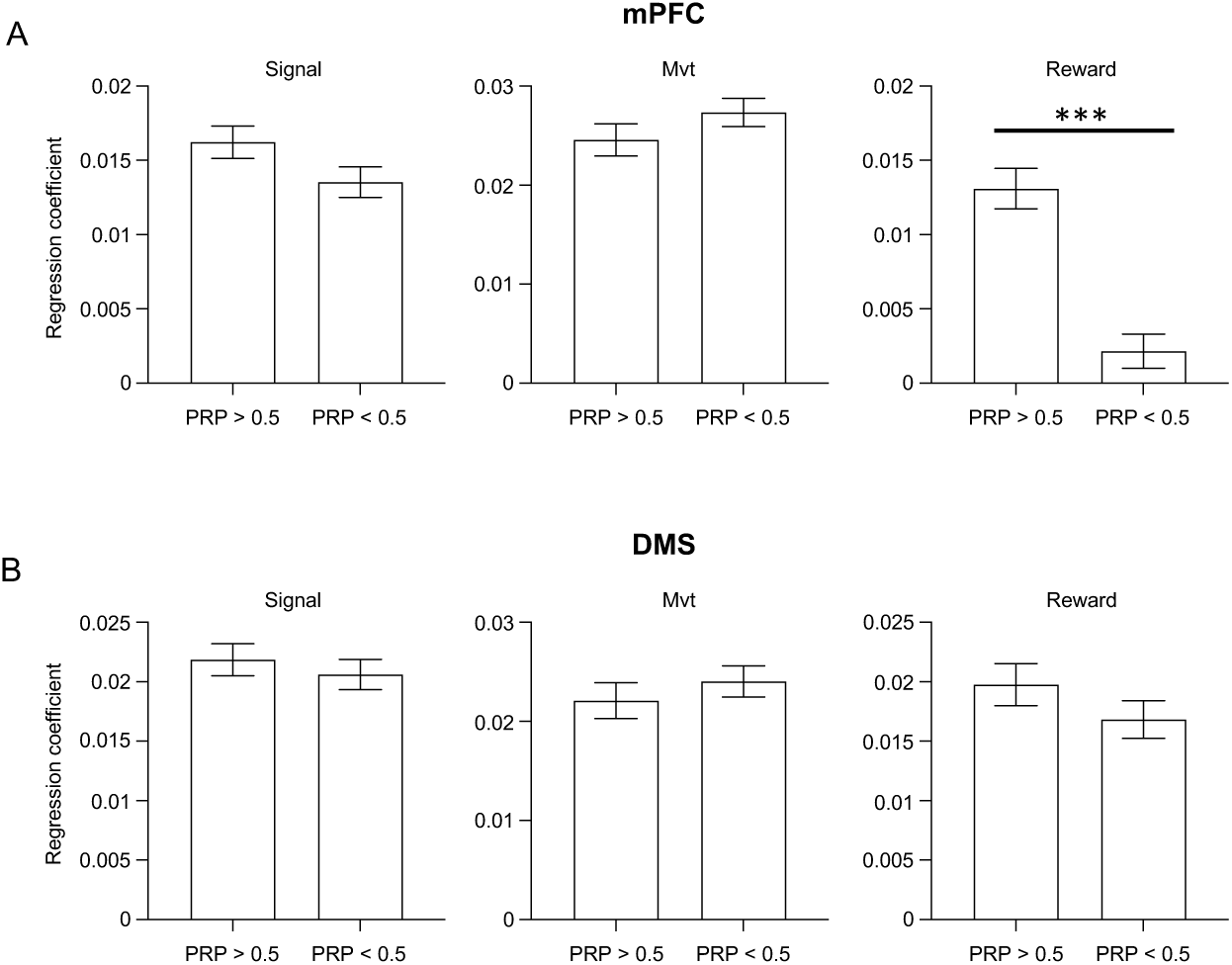
Overall reward history is represented in the mPFC but not in the DMS. **A.** mPFC regression coefficients for trials with low (<0.5) and high (>0.5) prior reward proportion (PRP – proportion of rewarded trials in the previous 10 trials). * indicates a statistically significant difference (5% confidence level, paired t-test). **B.** Same as G but for striatal neurons. Error bars indicate SEM.

### mPFC and DMS neurons encode the perceived reward probability difference which explains animal’s choice

Previous work showed a large number of mPFC neurons modulate with relative value (ΔQ) during both the intertrial interval (ITI) period and the task response window (TRW) period^15^. In agreement with this finding, Fig.5A shows raster and firing rate plots of an example mPFC neuron aligned by trial-initiation signal. When reward probability was high on the left and low on the right, and the PRPD was positive, its firing rate in both response period and intertrial period was high (and vice versa). We also encountered mPFC neurons that show an inhibitory response during each trial. The change in their firing rates can also modulate with reward probability. Fig.5B shows an example neuron whose firing rate during the response window was higher, thereby giving a smaller magnitude in its inhibitory response when PRPD was negative. Overall, we found that mPFC spike response magnitude during each trial can be positively or negatively correlated with PRPD (Fig.5E, left). There was no significant difference between the proportion of neurons that was significantly positively and significantly negatively correlated (Fig. 5E, right, proportions calculated for each session, paired t-test, P>0.05). Since PRPD is calculated by subtracting the ipsilateral reward probability from the contralateral reward probability this result indicates that the mPFC has no overall side bias. Interestingly, we also found the relative value or the PRPD was represented in the DMS as DMS neurons’ firing rates also modulated with PRPD in a similar fashion to mPFC neurons (Fig.5F, left). Fig.5C and D show two examples whose response magnitudes and intertrial firing rates modulate with changes in reward probability difference. DMS neurons also had no differences between the proportions that were positively or negatively correlated with PRPD (Fig.5F, proportions calculated for each session, paired t-test, P>0.05), suggesting that DMS has no side bias either.

**Figure 5.**
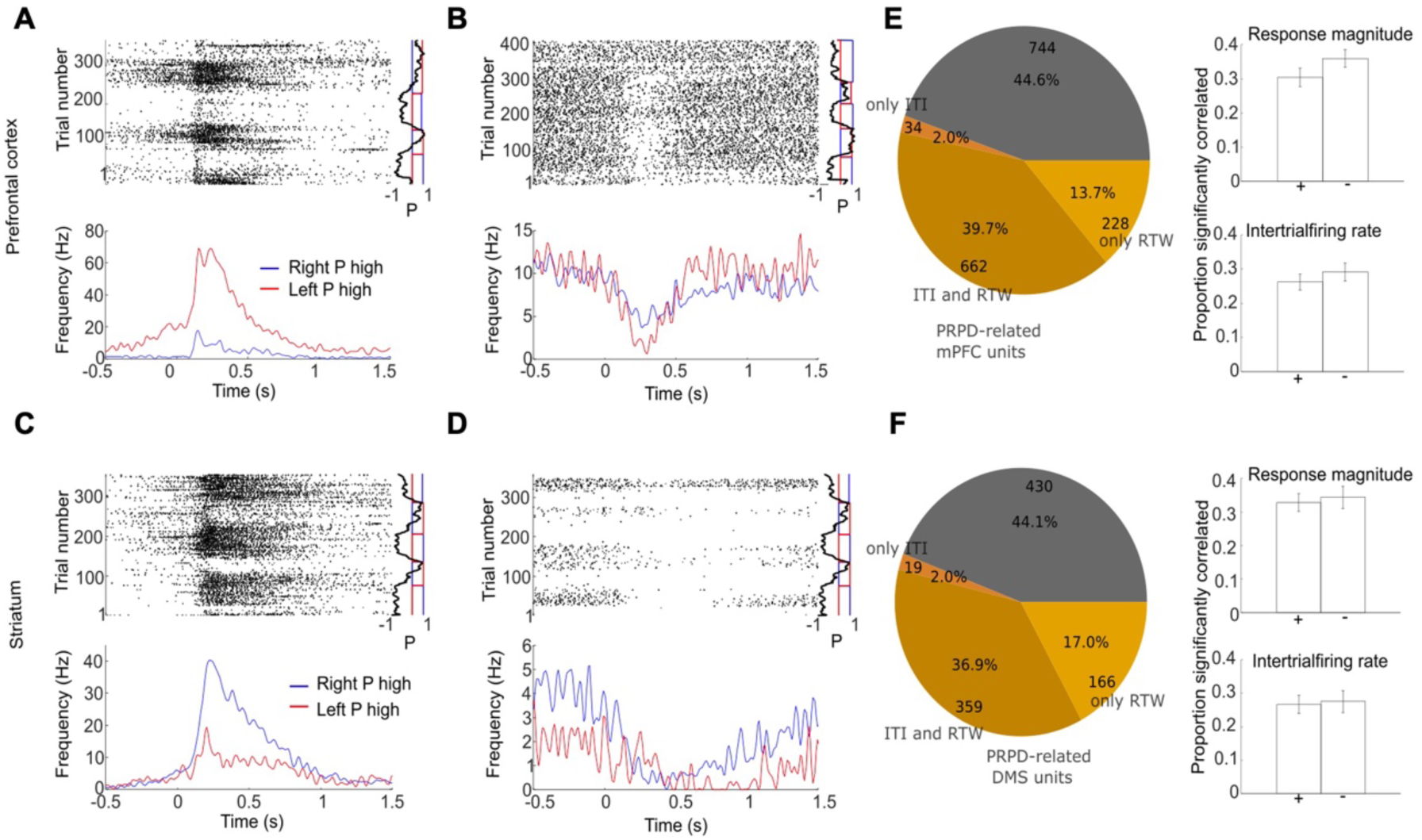
**Neural representation of PRPD that biases animals’ choices in mPFC and DMS**. **A-B**. Spike raster and firing rates of two example mPFC neurons. Firing rate plotted separately for high right reward probability (and low left) and high left reward probability (and low right). Vertical graphs plot the set probability for left and right (red and blue traces) and PRPD (black traces). **C-D.** Same plots as A and B, but for two example DMS neurons. **E.** Left: In all 1668 mPFC neurons, 924 (55.4%) were significantly related to PRPD when considering either response-time-window (RTW) or inter-trial interval (ITI) spikes. Right: Proportion of spiking response magnitudes and intertrial firing rates that are positively (+) and negatively (-) correlated with PRPD. Values were calculated for each session. **F**. Same analysis but for DMS neurons. In all 974 DMS units, 544 (55.9%) were significantly modulated by PRPD.

### mPFC spike modulation leads DMS spike modulation in response to changing perceived reward probability difference during dynamic foraging task

Considering PRPD is represented in both mPFC and DMS, given the strong anatomical connections between these two structures, we asked how mPFC neurons communicate with their DMS counterparts. To this end, we examined the temporal relationships between pairs of simultaneously recorded mPFC and DMS neurons over the course of the entire task session to gauge the inter-brain region dynamics of spike modulation during the foraging behavior. We first low-pass filtered the trial-wise intertrial firing rates (background firing) and the firing rate in the RTW (response magnitude) over the entire session (below 20 times the fundamental frequency of the session - see methods) and examined the phase difference between mPFC and DMS trial-wise spiking modulation. Fig. 6A and B show an example session with five switches and the trial-wise response magnitudes for an example mPFC-DMS cell pair over the entire session. Whilst the response magnitudes fluctuate from trial to trial, there is a clear slow and large fluctuation in both cells’ responses that mirror the PRPD plotted in panel A. The response magnitude traces were low pass filtered, and their phases were calculated via a Hilbert transform (Fig 6B, middle and bottom panel) showing a non-synchronous phase between the mPFC and DMS neurons. The circular-mean phase differences between each mPFC-DMS neuron pair were calculated for all sessions and their distribution was plotted in Fig.6C (top) for both response magnitudes and inter-trial frequencies.

**Figure 6.**
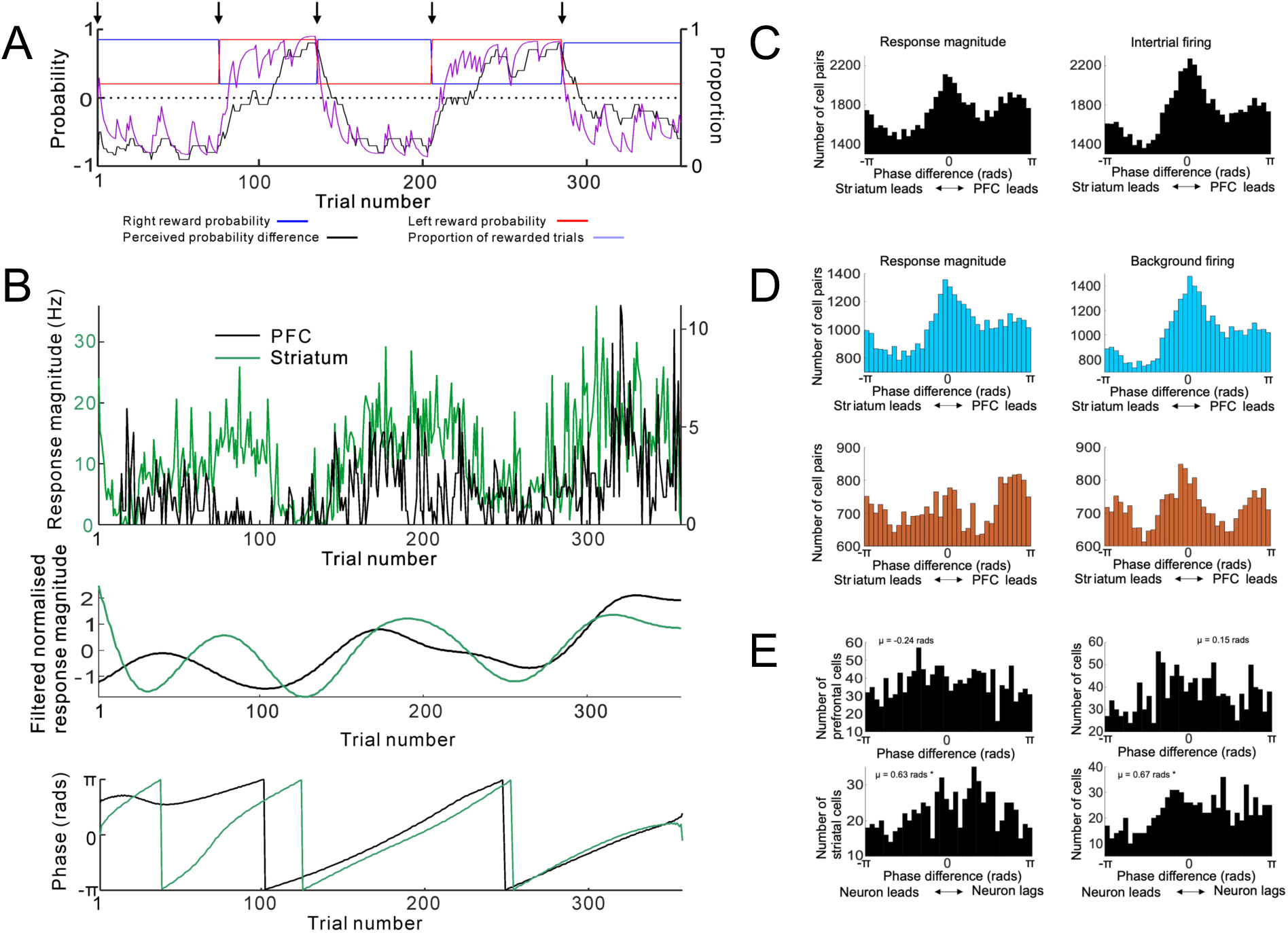
mPFC spike modulation leads DMS spike modulation in response to PRPD changes during dynamic foraging task. **A.**Set probability, reward proportion, and PRPD for an example session. Arrows indicate switches. **B.** Top: Example mPFC and DMS spiking response magnitude over the course of the same session. Middle: Low-pass filtered and de-meaned spiking response magnitude. Bottom: Trial-wise phases of the low-pass filtered spiking magnitudes. **C**. Distribution of mPFC-DMS phase differences of response magnitude and intertrial firing rate. Note significantly more cell pairs with phase differences between 0 and π. **D.** mPFC-DMS phase difference distributions as in C but separated according to task performance. Good performance sessions in blue, and poor performance sessions in orange. **E.** Distribution of phase difference between low-pass filtered PRPD and low-pass filtered neuronal spiking responses. µ indicates the circular-mean of phase differences. * indicates circular-mean to be significantly different to zero radians (P<0.05, circular m-test).

A mPFC and a DMS neuron that are both positively or both negatively correlated with PRPD will have a phase difference close to 0 whereas if one is positively correlated and the other is negatively correlated, then they will have a phase difference close to ±π. Since we have shown that roughly equal numbers of neurons are positively and negatively correlated with PRPD (see Fig. 5) one would assume that pair-wise mPFC-DMS phase differences would fall into a bimodal and symmetrical distribution with modes around 0 and ±π (i.e. roughly equal numbers of phase difference values between -π and 0 radians and between 0 and π radians). In reality, there are significantly more pairs with phase differences between 0 and π radians than between - π and 0 radians (P = 0.018 for response magnitude and P = 0.012 for intertrial firing rate, paired t-test; numbers of pairs calculated for each session), giving an asymmetrical appearance to the phase-difference distribution (Fig. 6C). This signifies an excess of cell pairs with a prefrontal lead, suggesting that overall mPFC neurons change their firing rates in response to the change of PRPD earlier than DMS neurons.

The example mPFC and DMS phases in Fig. 6B seem to have a greater variability relative to each other near the start of the session but a more consistent mPFC lead of DMS around the middle of the session and a more synchronous relationship near the end of the session. This could be due to an increasing familiarity with the rules of the task causing mPFC-DMS relationships to change from being fairly uncorrelated to a consistent mPFC lead and finally to mPFC-DMS synchrony. In other words, a good performance should be associated with a relatively more prominent peak about 0 radians and more positive values in its phase difference distribution (signifying more mPFC lead and more mPFC-DMS synchrony). Conversely, a bad performance should be associated with a more symmetrical phase difference distribution and a less prominent peak about 0 radians, and less positive values in phase difference distribution.

To test this hypothesis, we first gauged how well the animal followed the rules of the task for each session by calculating how well the response proportion to a given side followed the set probability for that side using a cross-correlation (normalized to give unitary autocorrelation). The cross-correlation for left and right is averaged for each session, and the peak value is used as a measure of the animal’s performance. Sessions were then separated according to whether their values were above or below average. The histograms of the mPFC-DMS phase differences are plotted in Fig. 6D for above- and below-average sessions (blue and orange). Above average sessions (good performance) have clear peaks around 0 radians (signifying prefrontal-striatal synchrony) as well as significantly more mPFC-DMS cell pairs with prefrontal lead (positive values in its phase difference), giving an asymmetrical phase distribution (significantly more phase difference values between 0 and π radians than between -π and 0 radians, P = 0.02 for response magnitude and P = 0.004 for intertrial firing rate, paired t-test; numbers of pairs calculated for each session). Below-average sessions (bad performance) have a largely symmetrical phase distribution without significantly more or fewer pairs showing mPFC lead or lag (P = 0.47 for response magnitude and P = 0.98 for intertrial firing rate).

We also take the mPFC and DMS neurons whose trial-wise spiking phases are significantly correlated with the phases of the PRPD (after similar filtering and Hilbert transform and calculating circular-to-circular correlation coefficients. For response magnitudes: 1346/1668 mPFC and 810/974 DMS neurons significantly correlated; for intertrial firing: 1340/1668 mPFC and 802/974 DMS neurons significantly correlated). We examined a possible phase difference between the PRPD and trial-wise spiking phases of these neurons in the mPFC and DMS respectively. The distributions of their phase differences are shown in Fig. 6E. The mPFC neurons’ phases are not significantly different from the PRPD phases (symmetric distribution of phase difference values with peak at 0), but DMS neurons show an overall lag (more positive values of phase difference, P<0.05, circular m-test of PRPD-neuron phase differences) – indicating that the majority of DMS neurons lag behind PRPD changes in their firing rate changes. Again, this agrees with an mPFC lead over DMS in firing rate modulation in the task.

### Putative monosynaptic pairs between the mPFC and DMS convey PRPD information

Given strong unidirectional excitatory connections between mPFC and DMS^30-32^, and the fact that a large proportion of neurons in the mPFC and DMS represent PRPD during our foraging task (see Fig.5), we examine the possibility of locating these representations anatomically at the level of monosynaptic mPFC-DMS pairs. To this end, putative monosynaptic excitatory (PMSE) pairs were sought between all possible mPFC and DMS pairs (Fig.7A). PMSE pairs were identified based on well-established methods with minor modifications^33,34^. The criterion is based on a strongly increased likelihood of spiking of the postsynaptic pair member with minimal jitter within a narrow time window (monosynaptic delay) after the presynaptic member’s spikes (see methods for details). Out of all 62774 possible mPFC-DMS pairs, 590 PMSE-pairs were identified. In roughly one-third of the PMSE pairs at least one member was related to PRPD, whereas in roughly one-fifth of them, both members encoded this decision variable (Fig.7B). These results provide direct functional evidence in support of mPFC-to-DMS transfer of PRPD information.

**Figure 7.**
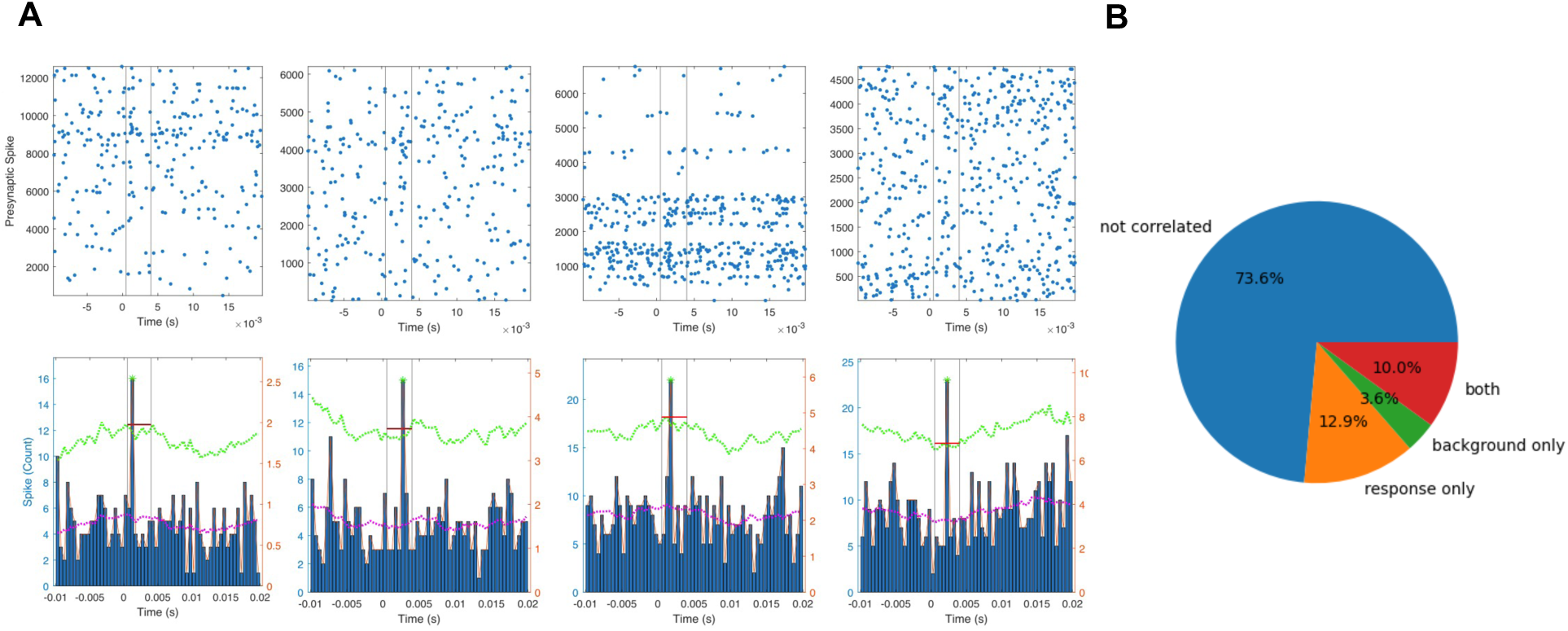
Putative monosynaptic pairs between mPFC and DMS convey PRPD information. Given the strong anatomical projections and PRPD encoding in both mPFC and DMS, putative monosynaptic excitatory pairs (PMSE; see Methods for details) were sought. 590 PMSE pairs were identified out of 62774 potential pairs (0.94%). **A.** Raster plots and cross-correlograms of 4 example PMSE pairs, with a narrow suprathreshold peak in the firing rate of the postsynaptic pair member within 0.5 and 7ms of the presynaptic member’s spikes. **B.** 26.4% of the 590 PMSE pairs, both members of the pair were significantly related to the PRPD; 12.9% were PRPD related based on response time window spikes only, 3.6% based on intertrial interval spikes only, while 10% were PRPD related based on both time windows.

### Lack of reward strengthens inter-region communication between mPFC and DMS during foraging

After elucidating the temporal relationship between the mPFC and DMS neural dynamics during foraging, we examined whether the strength of inter-brain region information transfer, e.g. to convey the decision signal PRPD, is modulated when the animal encounters a constant change of reward experience in the task. To test this, we calculated pair-wise spike-to-spike cross-correlations between mPFC and DMS neurons in the inter-trial periods (in order to avoid the effects of task-related co-modulations), normalized to negate the effects of changes in firing frequencies^35^. We use the absolute maximal deviation of spike-to-spike cross-correlation from zero to measure the strength of the interconnectivity (cell-to-cell interconnectivity) between a given mPFC-DMS cell pair. One example of this measure across an entire session (for inter-trial intervals spikes in a moving centered 20-trial long window) is plotted in Fig.8A along with prior reward proportion, PRP, concomitantly in Fig.8B. This example shows a significant negative correlation between interconnectivity strength and PRP (Fig.8C). Overall, a strong significant number of pairs have negative correlation between interconnectivity strength and PRP (Fig.8D), and the averaged cross-correlation across all pairs and all sessions was negative (Fig.8E) – indicating that periods of low reward experience lead to a strengthening of spike-to-spike transmission. Consistent with this interpretation, the cell-to-cell interconnectivity is significantly higher in non-rewarded trials than in rewarded trials across all pairs and all sessions (Fig. 8F). In addition, cell-to-cell interconnectivity is also significantly increased in trials during block transitions than in trials during block plateaus across all pairs and all sessions (Fig.8G).

**Figure 8.**
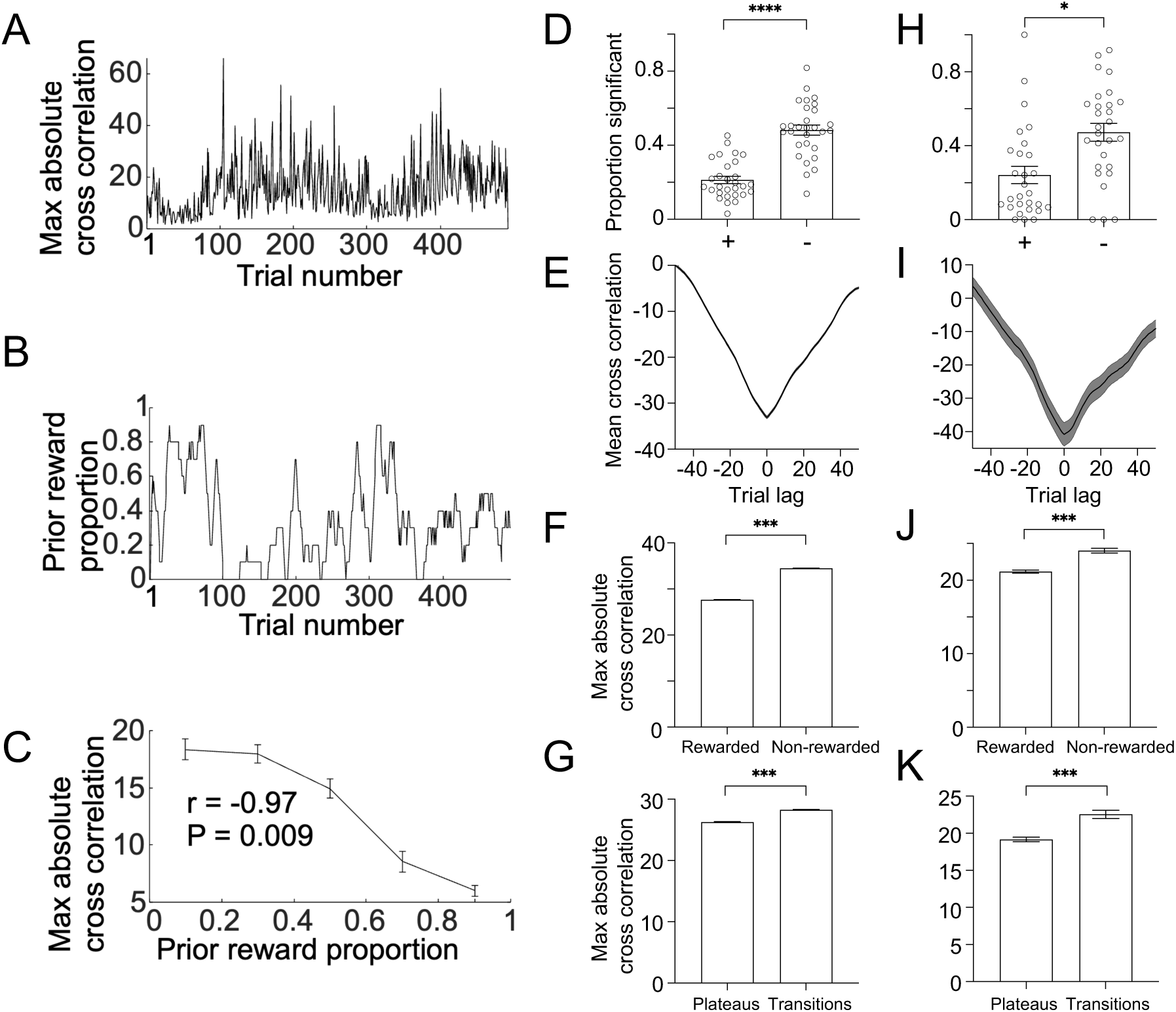
Lack of reward strengthens the connectivity between mPFC and DMS during foraging. **A-B.**The changes of maximum spike-to-spike cross-correlation strength of an example mPFC-DMS pair in A, and the change of overall prior reward proportion (PRP) in B, averaged over a centered moving 20-trial long window for an example session. **C.** The two signals from A and B have a strong negative correlation. **D.** The average proportion of negatively correlated pairs across all sessions was significantly higher than the positively correlated pairs. **E.** Mean cross-correlation between the degree of prefrontal-striatal spiking interconnectivity and reward proportion in all pairs and all sessions calculated for centered 20-trial-long moving windows. Cross-correlations normalized to give unitary auto-correlation at 0 lag. **F.** The average strength of interconnectivity (averaged maximal absolute cross-correlation) in non-rewarded trials was significantly higher than in rewarded trials. **G**. The average strength of interconnectivity was significantly higher in trials during block transitions (20 trials from the change of set probabilities) than in trials during block plateaus (20 trials before the change of set probabilities). **H-K**. Same analysis as D to G but for PMSE pairs, examining monosynaptic transmission fidelity between mPFC and DMS.

We also sought to examine the strength of any directly connected mPFC-DMS neuronal pairs^31^. To this end, we applied the above analysis in previously identified PMSE pairs between the mPFC and DMS as shown in Fig.7. In agreement with findings from all possible pairs, the spike-to-spike transmission fidelity between mPFC and DMS at the monosynaptic level was also enhanced when the animal experienced overall low reward proportions (Fig. 8H-K). Since PRP is a sensible measure of a need for behavioral change to an alternative advantageous strategy in our paradigm, these findings indicate that mPFC exerts enhanced influence over DMS when top-down control is beneficial.

## Discussion

In this study, we examined neural activities in the mPFC and DMS in head-restrained mice performing an action-outcome association based dynamic foraging task which requires internal tracking and updates of reward history to enable optimal performance in flexible decision-making. In summary, our results show: 1) neural population activity in the mPFC but not the DMS represents overall reward history information. 2) A large proportion of neurons in mPFC and DMS represent PRPD, and a quarter of putative monosynaptic mPFC-DMS pairs encode PRPD. 3) mPFC leads DMS in spiking modulation in response to PRPD changes and this temporal relationship is required for good performance. 4) Lack of reward strengthens the functional connectivity between mPFC and DMS.

The prefronto-striatal circuits are essential in flexible behavior which requires new learning, plasticity, and the implementation of chosen action plans, but the nature of their interaction is still not well understood. Two hypotheses were proposed, arguing for the leading role of stratum^19,20^ or prefrontal cortex^21,22^, respectively, in detecting and representing circumstance changes in flexible behavior. Our results support the second hypothesis where mPFC is in the center as a ‘context detector’ and trains striatum. More recently, an updated, reconciling hypothesis from non-human primate studies postulates that DMS representation will appear early in flexible behavior involving pure stimulus-response learning^36^, whereas when cognitive demand is higher, e.g. in tasks requiring the learning of abstract or inferred information such as rules and categories, prefrontal cortex will encode such changes earlier^37,38^. Our results here provide direct evidence to support this theory and expand this hypothesis to rodents. The fact that mPFC leads DMS and overall reward history information is held only in mPFC suggests that in a response-outcome association where the subjects have to rely not on learning perceptual information (visual or auditory stimuli) per se but self-inferred concepts such as reward history, the mPFC will play a leading role in this circuitry. In addition, our data suggest that the establishment of this mPFC-leads-DMS relationship is essential for successfully executing this form of flexible behavior, as evidenced by the fact that when this temporal coordination is lost task performance is impaired and vice versa. Our results further demonstrated that striatal activities not only faithfully reflect sensorimotor activities in corresponding cortical regions^39^, but also inherit the representations of more abstract and cognitively inferred information, from high-order associative cortices such as the mPFC.

The dynamics of functional connectivity between mPFC and DMS in flexible behavior is poorly understood, likely due to limited studies involving simultaneous prefrontal and striatal recordings in task-performing animals that switch responses or behavioral strategies back and forth several times in a single session, involving periods of learning, learned, and new learning several times in one experimental episode. Our experimental design provides such an opportunity. By employing simultaneous double silicon probe recordings in two separate brain regions during behavior, we have identified putative monosynaptic mPFC-DMS pairs that are jointly modulated by decision variables of the task. To our knowledge, this is the first attempt to delineate neural dynamics in the prefronto-striatal connectivity at the level of real-time, cross-region spike transmission. Our results provide direct functional evidence at the cellular level with defined anatomical specificity to how mPFC and DMS cooperate in real-time in implementing flexible behavior. Our results show that the strength of information transmission between mPFC and DMS during flexible behavior is not static but dynamic, even at the level of monosynaptic pairs. It is negatively modulated by the overall prior reward proportion, a behavioral variable likely associated with an animal’s overall drive and vigor^15^. How is this modulation achieved in the corticostriatal circuit? The reward prediction error theory postulates that a teaching signal conveyed by dopamine is critical for multiple forms of associative learning by modulating corticostriatal synaptic plasticity in striatal medium spiny neurons^40^. Previously, reward prediction errors were commonly observed in behavioral paradigms of explicit stimulus-outcome and stimulus-response associations^41,42^. The fact that lack of reward enhances functional connectivity between mPFC and DMS in our experiments may be explained by a recent finding of action-outcome (response-outcome) prediction error signals in goal-directed behavior, conveyed by nigrostriatal dopamine projections^43^. In such a scenario, dopamine release is selectively suppressed in the reinforced actions associated with the expected outcome (advantageous choices in trials in the plateau period in our experiment) to minimize new learnings in ‘learned periods’ for predicted outcomes (expected rewards), permitting performance to stabilize. During the block transitions, this suppression of dopamine in previously reinforced actions may facilitate the standing-out of unexpected reward signals from unreinforced actions and/or error signals following reward omission^16^ which promotes new learning in pursuing alternative choices. Nevertheless, uncovering anatomically defined circuit mechanisms responsible for this response-outcome reward prediction error which may underlie the observed strengthening of corticostriatal transmission efficacy by lack of reward during flexible behavior remains an important question for future studies.

In conclusion, our results raise the feasibility of a framework where the mPFC tracks the action-outcome history faster and earlier than the DMS in flexible behavior. The computed decision signal is then sent down to the DMS via deep-layer corticostriatal projection neurons to bias behavioral engagement toward an advantageous strategy. In such a model, DMS or basal ganglia can be considered as a behavioral-output filter, representing the engaged strategy, which can be fast set and reset by mPFC when there is a need for a change. mPFC senses the need for change probably via a circuit mechanism that may involve dopamine-mediated response-outcome reward prediction error signal and feedback monitoring^44^. Then, when mPFC-to-DMS functional connectivity is enhanced, high-order influence from mPFC will be able to liberate basal ganglia’s engagement from a chosen strategy and re-assign the basal ganglia to a new and advantageous one, thus enabling a fast behavioral adaptation in response to changing environmental variables.

## Material and methods

### Surgical preparation

All animal procedures received prior written approval from the UK Home Office and local ethical and animal warfare committee under the project license PP8564759.

3-5 months old male C57/BL6J mice (Charles River, UK) were anesthetized with isoflurane (4% induction, 1-2% maintenance) and mounted into a stereotaxic frame. The skull was surgically exposed, cleaned, and covered with a layer of cyanoacrylate superglue (Loctite, Germany). The sites of craniotomies were marked. Two steel screws (1.3 mm diameter) were placed in small boreholes drilled above the cerebellum and olfactory bulb. A custom-made head-ring for later head-restraint training and recording was attached to the skull and the screws and tightly fixed using bone cement (Refobacin, Biomet). After the surgery, mice were injected with Carprofen (Rimadyl,

120-130 µL S.C., 20 mg/kg) and received a liquid supplement with 0.9% NaCl, 200-300 µL I.P.. Mice were given at least 7 days to recover with health monitored. Mice were then handled and habituated to be head-restrained on a custom-modified wheel-turning apparatus^25^ adapted into a virtual reality (VR) system (Phenosys, Germany). After habituation, mice were water-restricted (approximately 1.2ml water/day) and trained to perform the foraging task by turning the wheel for liquid rewards. Daily training lasted about 30-40 minutes. When the animals were deemed to have learned the dynamic foraging task (successful 4-5 block transitions in one training session on two consecutive days) after 3-6 weeks, they were anesthetized, and two small craniotomies were drilled. One above the medial prefrontal cortex (AP: Average: 2.1mm; Range:1.8- 2.35mm; ML: Average:0.71mm; Range: 0.01-1.01mm) and one above the ipsilateral (4 mice) or contralateral (1 mouse) dorsomedial striatum. The craniotomy positions were adjusted to the applied silicon probe angles, for the DMS probe on average 14.5° antero-posterior angle, and for the PFC probe on average 15.3° medio-lateral angle was used (right hemisphere 4 mice, left hemisphere 1 mouse). Dura was removed. Craniotomies were sealed and protected by silicone (Kwik-Cast, World Precision Instruments, USA). Animals were left for recovery for at least 8 hours before electrophysiological recordings started.

### Action-outcome based dynamic foraging task

Animals were head-fixed within a 180-degree virtual-reality screen (PhenoSys, Germany). Their front paws were placed on a wheel which they were able to move counter- and clockwise (left or right, see Fig.1A). This moves an on-screen cursor left or right. Fluid rewards were delivered in volumes of 4µL per rewarded trial via a tube placed in front of the animal’s mouth. Each trial starts with the appearance of the cursor in the centre of the screen and a trial initiation signal (auditory tone of 8000Hz lasting 0.03s). After the animal rotates the wheel, the reward is delivered in a probabilistic fashion. One side will have a high reward probability (discrete values between 0.7 to 0.9), and the other side will have a low reward probability (discrete values between 0.05 to 0.2). After a variable number of trials, the high and low reward probabilities switch sides, so that the previously advantageous side becomes disadvantageous. Block transition was based on multiple conditioning. When the trial number reaches a randomly selected discrete value between 50 and 110, the animal’s performance is checked. If it is over the 75% advantageous choice ratio, a block transition is triggered. If it does not, a randomly selected number of extra trials between 15 and 30 are added to the current block, at the end of which performance is rechecked. After the animal executed wheel-turning (and a reward is either delivered or not delivered), there was a pause of variable duration (inter-trial interval, ranging from 1s to 1.6s after responded trials and 1.9s to 2.3s after non-responded trials). The variable duration is to make it more difficult for the animals to anticipate when the next trial starts, and thus discourage rhythmic premature responding and encourage attentive responding after the trial initiation signal. After the end of the variable pause period there is a period lasting 1s when the animal is not permitted to make any suprathreshold movements of the wheel. If such movements occur, the time window resets to the start of the 1s. At the end of this period, the next trial commences. If the animal makes no suprathreshold movement for 7s after the trial initiation signal, a new trial is started. The times of the trial initiation signals and rewards were recorded as digital pulses simultaneously with neuronal signals via the signal-acquisition device’s digital inputs. The wheel’s instantaneous angular velocity was calculated from the raw outputs of its 1024-tick quadrature rotary encoder, also simultaneously recorded.

The proportion of rewarded trials and the proportion of responses towards the high probability side (advantageous responses) were calculated using a centered moving 20- trial long window. Perceived reward probability was gauged by calculating, for each side, the proportion of rewarded trials in the previous 10 trials of responses made to that side.

### In vivo electrophysiological recordings and signal processing

Electrodes consisting of 128-contacts on 4 parallel shanks (A4X32-Poly2-5mm-23s- 200-177, NeuroNexus, USA) were inserted into the medial prefrontal cortex (mPFC) and dorso-medial striatum (DMS). Each shank contained 2 columns of contacts with horizontal and vertical inter-contact distances of 30 and 36µm. Contact impedance values ranged from 0.5 to 3MΩ. Signals were recorded using an Intan 128-channel head-stage from each electrode and an Intan RHD recording controller. Signals were sampled at 20KHz and digitized as 16bit signed integers. Spikes were discriminated using the software Kilosort version 2.0^45^. Automatically clustered waveforms were post hoc manually curated using the software Phy (https://phy.readthedocs.io/en/latest/). Signal processing and data analysis were carried out in Matlab (Mathworks, USA) or Python. All filtering in the study was carried out using a 4-pole Butterworth filter filtering in both directions.

### Multiple linear regression analysis of task modality representation in spike firing profile

Cell firing rate profiles are aligned by trial initiation signals (−0.5 to 1.5s) using 20ms-wide bins and then low-pass filtered at 20Hz. In order to gauge the degree of representation of different task events in spiking profile we first separated trials into: 1) those that had only the trial initiation signal but no subsequent suprathreshold movements or rewards (S trials); 2) those with trial initiation signal and movement but no reward (SM trials); 3) those with signal, movement and reward (SMR trials). (For all recorded neurons from all sessions). In order to derive spike firing modulations that represent the trial initiation signal we simply took the average firing rate profile for S trials (see Fig.3B and C). To derive movement-related firing modulations we subtracted spiking profiles of S trials from SM trials and averaged across all neurons in each area. In order to derive the reward-associated spiking modulation we subtracted the spiking profiles of SM trials from SMR trials. We used these three modality-related modulation patterns (S,M and R, representing signal, movement, and reward, scaled to have magnitudes of 1) as basis functions with which to fit individual neuronal firing profiles via a multiple linear regression:

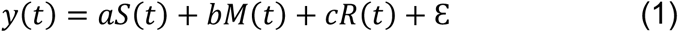

where y is the modulation of an individual cell’s firing rate in SMR trials (i.e. where all three task modalities are present). S, M and R are the modality-associated basis functions for signal, movement and reward; a, b and c are their least-squares fitted coefficients; Ɛ represents the overall error of the regression fit. The values of a, b, c, and K are derived with the aim of producing the smallest value of Ɛ.

In order to guard against over-fitting, we carried out a 10-fold cross-validation. We randomly sampled 10% of neurons in each area as the training set and used their firing profiles in SMR trials to generate the modality-associated basis functions and their respective coefficients. Using these values, we generated a fitted firing profile and compared it with the actual firing profiles of the remaining 90% of neurons via a Pearson’s correlation analysis. This was repeated 10 times, each time with a distinct set of neurons in the training set. The correlation coefficients for all iterations are all significantly positive (P<<0.001, one-sample t-test).

### Phase calculation of trial-wise neuronal firing modulations

In order to examine broad and slow changes in the trial-wise spiking responses of neurons throughout the entire recording session we calculated for each trial the intertrial firing rate and response magnitude. Intertrial firing rate was defined as firing frequency for times between −1s and −0.5s relative to trial initiation signal. Response magnitude was defined as the mean absolute difference between firing rate in the 0 to 1.5s window and mean intertrial firing rate. We then low-pass filtered these signals using a cut-off frequency that is 10 times the fundamental frequency of the session (the fundamental trial frequency of each session is simply 1 divided by the number of trials), using a Butterworth filter. This allowed us to capture any oscillatory spiking process with frequencies up to 10 cycles per session. To calculate the trial-wise phase of the filtered signal we removed its mean using detrend function and performed a Hilbert transform.

### Spike-to-spike cross correlation

As a gauge of the degree of overall interconnectivity between pairs of prefrontal and striatal neurons we binned their spike times in 10ms-wide bins and then performed a normalized cross correlation calculation. The normalization procedure expresses the cross-correlation as the proportional difference of the raw cross-correlation relative to expected cross-correlation for uniform firing with the same mean rate as the actual neurons ^46^. This approach is designed to negate the effects of differences in spike firing rate on cross correlation. The normalized cross correlation *x́*(*τ*) between two time series *g*(*t*) and ℎ(*t*) is calculated using:

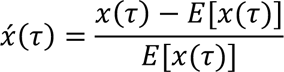

Where:

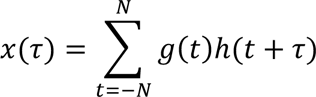

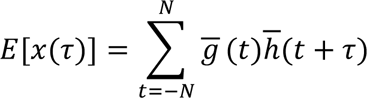

where *g*(*t*) and ℎ(*t*) are the binned spike firing for two spike trains, *x*(*τ*) is their raw cross correlation at lag *τ*, *t* represents time point, *N* represents maximal lag. *E*[*x*(*τ*)] is their expected cross-correlation and 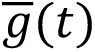 is a time series with equal length to *g*(*t*), where each time point equals the mean of *g*(*t*). The same applies to 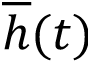.

Because the spike-pair correlations can be both positive and negative, the degree of interconnectivity between a given pair of neurons is defined as its maximal absolute deviation from zero. To adjust for the sudden variations in the firing rates, the interconnectivity strength for each trial is calculated using a centered 20-trial long window. In order to remove the effects of task-driven co-modulation on spike cross correlation we only used spikes in the inter-trial interval.

### Detection of putative monosynaptic pairs in the mPFC and DMS

Putative monosynaptic excitatory (PMSE) pairs were identified based on an increased likelihood of spiking of the postsynaptic pair member in a short latency, narrow time-window after the presynaptic member’s spikes in a cross-correlograms based analysis modified from previously published methods^33,34^. Cross-correlograms were constructed as postsynaptic-member spike count histograms, from −10 to +20ms time windows around every presynaptic spike, in 0.5ms bins. All pairs which fulfilled the following 4 conditions, were identified as putative PMSE pairs. I.) The postsynaptic member spike count was 3-SD above the expected spike count threshold in a 0.5 - 7ms time window after the presynaptic spikes. For the cortico-striatal pairs a longer delay time window was selected compared to what is usually used in local circuits where the synaptic delay is short, with EPSP latencies in the range of 0.5 to 3.5 ms^47,48^. This is also reflected in previous studies with a maximum delay for identifying PMSE pairs in cross-correlograms (up to 5 ms) in the hippocampus^33,34,49^. In cross-region cortico-striatal projections, actional potential conduction velocity is slower than the pyramidal tract and other faster pathways^50^, resulting in a peak probability of postsynaptic spiking at 5-7ms delay but frequently up to 11ms^51^. Therefore, our 0.5-7ms delay time window for identifying PMSE pairs was chosen accordingly. The expected spike count for every bin of each pair was then calculated based on shuffling postsynaptic spike times with a ±5 ms jitter for 500 times. II.) There were at least 10 spikes above the expected line in the peak. III.) The ratio of spikes above the expected count and the expected count itself is above 1.5. IV.) The peak width was not longer than 1 ms (FWHM??), as low jitter is an indication of a monosynaptic interaction to guard against polysynaptic interactions.

### Reinforcement Learning Modeling

In addition to the intuitive PRPD, reinforcement learning modeling was also utilized in to quantitatively describe the animal’s foraging behavior. The Rescorla Wagner model commonly used in non-stationary two-arm bandit tasks was chosen to be a suitable candidate.

The model (sometimes referred to as the Q-learning model in other publications^15,16^)’s decision is guided by two decision variables, associated with two alternatives given to the animals. In our case, *Q*_*l*_ corresponds to leftward choices, while *Q*_*r*_ corresponds to the right. They were fed into the softmax function, along with a bias term *b* and inverse temperature parameter *β* to produce the probability of making a rightward choice.

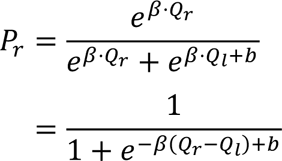

The second equation is the result of dividing both the numerator and denominator by *e*^β⋅Qr^ and is used in the optimization process.

After each trial, the model updates the decision variables using the following equations:

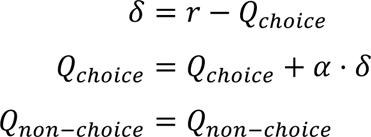

Where *δ* is the prediction error, *α* is the learning rate, *r* is the reward received, *Q*_choice_ is the decision variable associated with the chosen side, *Q*_non-choice_ is the decision variable associated with the side not chosen by the animals.

The optimization process aims to minimize the negative log-likelihood of the model’s choice probability given the actual choices made by the animals. The negative log-likelihood is calculated using the following equation:

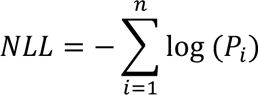

Where *n* is the number of trials, *P*_*i*_ is the probability of making the animal’s choice at trial *i*. The optimization process is handled by Python’s *Scipy* package, using the Nelder-Mead method of the ‘minimize’ function. The optimization process was repeated 100 times for each session, with the initial parameters randomly sampled from a uniform distribution to avoid local minima as much as possible.

The parameters were bounded to the following ranges during the optimization process to guarantee a meaningful result:

|  |  |  |
| --- | --- | --- |
| $\beta$ | 0.1 | ∞ |
| $\alpha$ | 0.001 | 1 |
| $b$ | $-\infty$ | ∞ |

